# Contemplative mental training reduces hair glucocorticoid levels in a randomized clinical trial

**DOI:** 10.1101/2020.11.13.381038

**Authors:** Lara M.C. Puhlmann, Pascal Vrtička, Roman Linz, Tobias Stalder, Clemens Kirschbaum, Veronika Engert, Tania Singer

## Abstract

**Objective:** This study had the objective to investigate the effect of regular contemplative mental training on endocrine indices of long-term stress.

**Methods:** An open-label efficacy trial comprising three distinct 3-month modules targeting attention and interoception, socio-affective or socio-cognitive abilities through dyadic exercises and secularised meditation practices was designed and carried out in 332 healthy meditation-naive adults. Participants underwent the training for up to 9 months or were assigned to a retest control cohort. Chronic stress indices were assayed at four timepoints, i.e., pre-training and following each module. The main outcome measures were cortisol and cortisone concentration in hair and self-reported chronic stress

**Results:** N=362 initial individuals were randomized, of whom n=30 dropped out before study initiation, n=4 before first sampling and n=2 were excluded. N=99 participants did not provide hair samples. Data from three separate training cohorts revealed consistent decreases in hair cortisol and cortisone levels over the training period. This effect increased with practice frequency, was independent of training content and not associated with change in self-reported chronic stress.

**Conclusions:** Our results point to the reduction of long-term cortisol exposure as a mechanism via which contemplative mental training may exert positive effects on practitioners’ health.

**Trial registration:** ClinicalTrials.gov identifier: NCT01833104

## 1. Introduction

Rising prevalence of stress-related mental and physical disorders (Habib & Saha, 2010; Vos et al., 2015) has led to the recognition of chronic stress as one of the 21^st^ century’s major health risks (Rosch, 2001). The health outcomes of exposure to psychosocial stress are mediated by prolonged activation of our main endocrine stress systems, the sympathetic-adrenal-medullary (SAM) and the hypothalamic-pituitary-adrenal (HPA) axes. Both systems exert complex effects on immune and metabolic processes and are causally involved in the development of cardiovascular, metabolic, and autoimmune disorders, among others (Chrousos, 2009). In striving to reduce stress and promote health and wellbeing, secular meditation-based mental training interventions, such as the mindfulness-based stress reduction (MBSR) program (Kabat-Zinn, 1994), have gained popularity. Various health-related benefits have been associated with engagement in such training interventions (see e.g. Grossman et al., 2004; Khoury et al., 2015 for meta analyses).Findings from our own 9-month mental training study, the ReSource Project (Singer et al., 2016), show differential positive changes in subjective well-being, cognition, peripheral physiology, and brain plasticity following distinct types of contemplative mental training (Singer & Engert, 2019).

Of particular interest for clinical application are the downstream health benefits of contemplative training, such as mitigation or prevention of stress-related disorders. Current theory suggests that these outcomes are mediated by dampened activity of physiological stress systems, above all the HPA axis (e.g. Creswell & Lindsay, 2014). In line with this theory, subjective-psychological stress load is one of the most widely reported training outcomes (e.g. Khoury et al., 2015). At the same time, self-report measures of contemplative training effects may be particularly vulnerable to confounds such as demand-effects and expectancy bias, since the training trials are inevitably open-label. Researchers are thus increasingly relying on physiological measures as more reliable and objective health outcomes. Results from these studies have shown that although correspondence between psychological and physiological measures of stress is often assumed, evidence for training-related endocrine stress reduction in healthy participants is currently mixed and inconclusive.

Studies of mental training effects on stress-related biomarkers predominantly focus on secretion of the HPA axis output hormone cortisol, either in response to acute stress or during basal activity, measured in blood or saliva. We previously found reduced cortisol secretion in response to an acute psychosocial laboratory stressor following the 3-month long training of socio-affective or socio-cognitive practices, but not after the training of present-moment attention and interoception (Engert et al., 2017). Several other studies of psychosocial stress induction found no effects of mindfulness- or compassion-based training on acute cortisol release (Arch et al., 2014; Rosenkranz et al., 2013; for a review see also Morton et al., 2020). Similarly heterogeneous results emerge at the level of basal HPA axis activity. Reports of lower diurnal cortisol output, mainly following mindfulness-based interventions employing the MBSR program (e.g. Brand et al., 2012), are contrasted by several null results (for meta analyses, see Pascoe et al., 2017; Sanada et al., 2016). These mixed outcomes do not satisfactorily corroborate the hypothesis that reduced HPA-axis activity mediates long-term training-related health benefits. Notably, however, while acute and diurnal cortisol indices provide a window to an individual’s long-term cortisol exposure, both bear shortcomings as measures of chronic stress. Cortisol levels collected after acute challenge reflect stress responses in a highly specific setting, and indices of diurnal cortisol measured in saliva, blood, or urine fluctuate considerably from day-to-day (Law et al., 2013; Ross et al., 2014). Since it is the long-term, cumulative HPA axis activation that is particularly maladaptive and related to ill-health (Chrousos, 2009; McEwen, 2000), methodological limitations in capturing chronic physiological stress may account for some of the heterogeneity in the contemplative training literature.

The present study aimed to investigate whether contemplative mental training affects patterns of long-term cortisol secretion as a mediator of downstream health benefits in 227 healthy adults. Instead of acute or diurnal cortisol secretion, we utilized the method of hair cortisol (HC) and cortisone (HE) assessment as indices of physiological long-term stress. HC and HE concentrations are assumed to capture systemic (i.e., whole body) cortisol exposure and have been linked to the subjective experience of psychosocial stress (Stalder et al., 2017). HC concentration is also positively correlated with diurnal cortisol output (Engert et al., 2018; Stalder et al., 2017) but less prone to state-related variance, which may allow for a particularly stable prediction of whether mental training has a long-term impact on HPA axis activity. Alongside cortisol, it has been suggested that levels of the inactive cortisol metabolite and precursor molecule cortisone yield a complementary, potentially more stable glucocorticoid signal (Stalder et al., 2013, supplement). We thus assayed cortisol and cortisone levels in 3 cm proximal hair segments, corresponding to approximately 3 months exposure. Additionally, self-reported chronic stress was measured using the Perceived Stress Scale (PSS; Cohen, Kamarck, & Mermelstein, 1983) and the Trier Inventory for Chronic Stress (TICS; Schulz & Schlotz, 1999).

The training regimen of the ReSource Project was designed to examine the specific effects of three different types of mental practice on HC and HE levels. This differentiated approach is especially valuable given the multifaceted nature of many mindfulness-based programs, which typically combine diverse practice types (Dahl, Lutz & Davidson, 2015). In three separate modules termed Presence, Affect, and Perspective participants trained attention-, socio-emotional, or socio-cognitive based practices, respectively, for 3-months each (Figure 1A). Participants were assigned either to one of two 9-month training cohorts completing all three training modules in different orders (TC1 and TC2), a 3month Affect only training cohort (TC3) or a retest control cohort (RCC) (Figure 1B; Singer et al., 2016, chapter 7). During each module, participants completed a standardized training routine involving weekly 2-hour group sessions and daily online practice. When examining effects on cumulative, slow changing indices like HC and HE, such rigorous, longitudinal training is arguably most promising.

**Figure 1.**
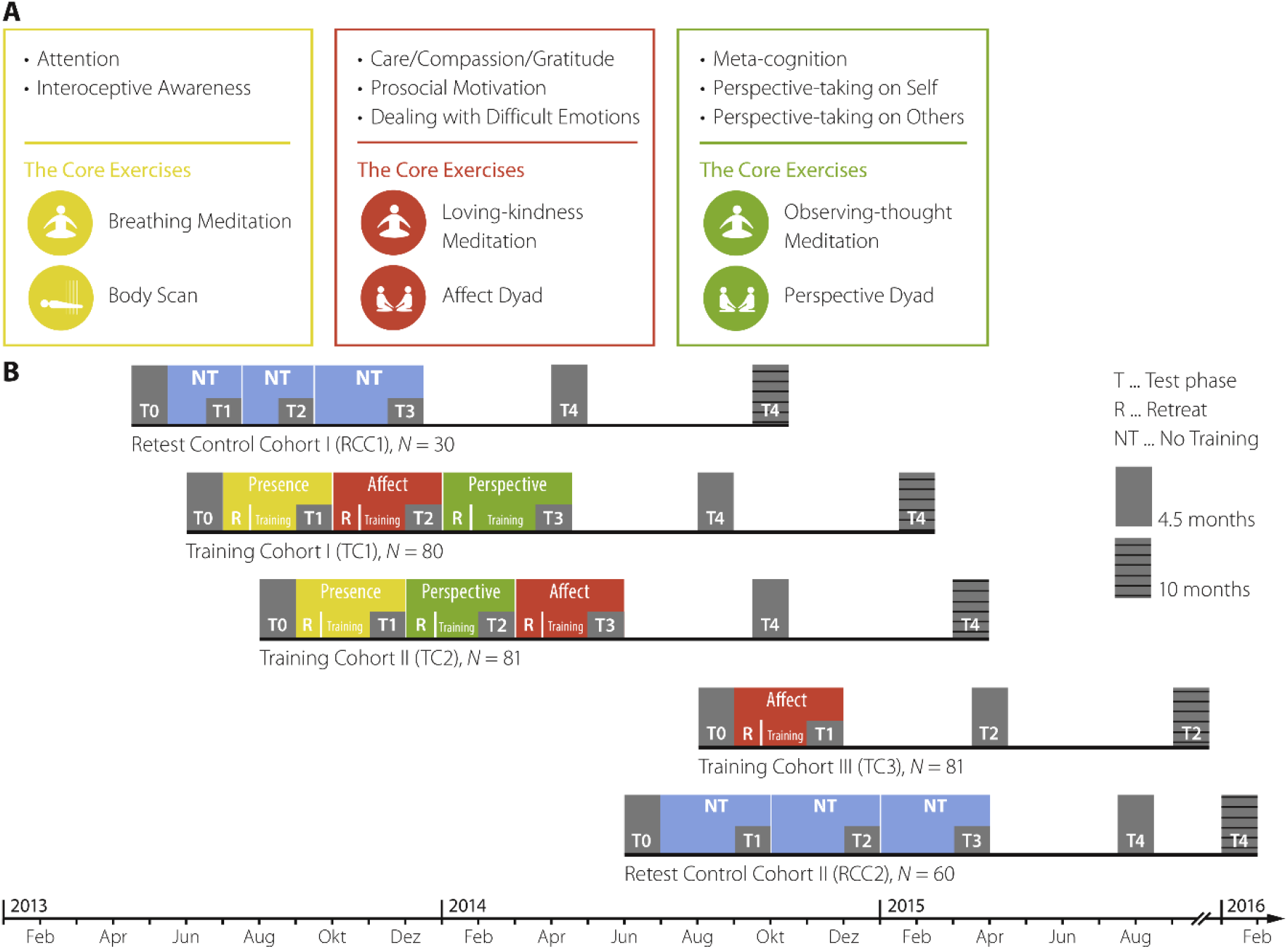
Study protocol and design. A) Core processes and practices of the ReSource training. The Presence module aims to train attention and interoceptive body awareness; its two core practices are Breathing Meditation and Body Scan. The Affect module targets social emotions such as compassion, loving kindness, and gratitude; core practices are Loving-kindness Meditation and Affect Dyad. In the Perspective module, metacognition and perspective-taking on self and others are trained through the core practices Observing-thoughts Meditation and Perspective Dyad. B) Design and timeline of the Resource Project. Two training cohorts TC1 and TC2 started their training with the mindful attention-based Presence module. They then underwent the social Affect and Perspective modules in different orders. The total training time for TC1 and TC2 was 39 weeks (13 weeks per module). TC3 only trained the Affect module for 13 weeks and the two RCC completed all the testing without training (for more detailed information see chapter 4 in Singer et al., 2016). Figure adapted from Singer et al. (2016), chapter 3.

In light of the above outlined evidence for changes in diurnal cortisol after mindfulness-based training, we primarily expected to find a reduction in HC and HE levels after the attention-based Presence module, which included classic mindfulness-practices that are also central to the MBSR program. Because basal and stress-induced cortisol levels are not reliably associated (e.g. Kidd et al., 2014), it remained an open question whether the acute stress-reducing properties of the social Affect and Perspective modules identified in our previous study (Engert et al., 2017) would translate to reduced cortisol levels in hair.

## 2. Materials and Methods

### 2.1 Participants

All participants underwent comprehensive face-to-face mental health diagnostic interviews with a trained clinical psychologist and completed additional mental health questionnaires. Volunteers were excluded if they fulfilled the criteria for an Axis-I disorder within the past two years or for schizophrenia, psychotic disorder, bipolar disorder, substance dependency or any Axis-II disorder at any time in their life. Volunteers taking medication influencing the HPA axis were also excluded (for further details on the screening procedure, see Singer et al., 2016, chapter 7). The ReSource Project was registered with the Protocol Registration System of ClinicalTrial.gov (Identifier NCT01833104) and approved by the Research Ethics Boards of Leipzig University (ethic number: 376/12-ff) and Humboldt University Berlin (ethic numbers: 2013-20, 2013-29, 2014-10). The study was conducted in accordance with the Declaration of Helsinki. Participants gave written informed consent, could withdraw from the study at any time and were financially compensated.

To avoid straining participants through excessive testing in the context of the multi-measure ReSource Project, sampling of hair was presented to participants as an optional rather than a core testing procedure, leading to lower adherence rates. Of 332 initial ReSource participants (197 women; mean age ± SD: 40.74±9.24 years; age range: 20-55 years), 217 provided hair samples at baseline (T0), of which 179 could be re-assayed for the present change analysis; 157 provided samples at T1, 136 at T2 and 150 at T3 (see Figure 2 and Tables S1 and S2 for sample sizes of all measures per cohort and reasons for missing cases). Twenty-four participants (18 women) were light smokers (≤10 cigarettes/day; mean ± SD: 16.01±16.09 cigarettes/week).

**Figure 2.**
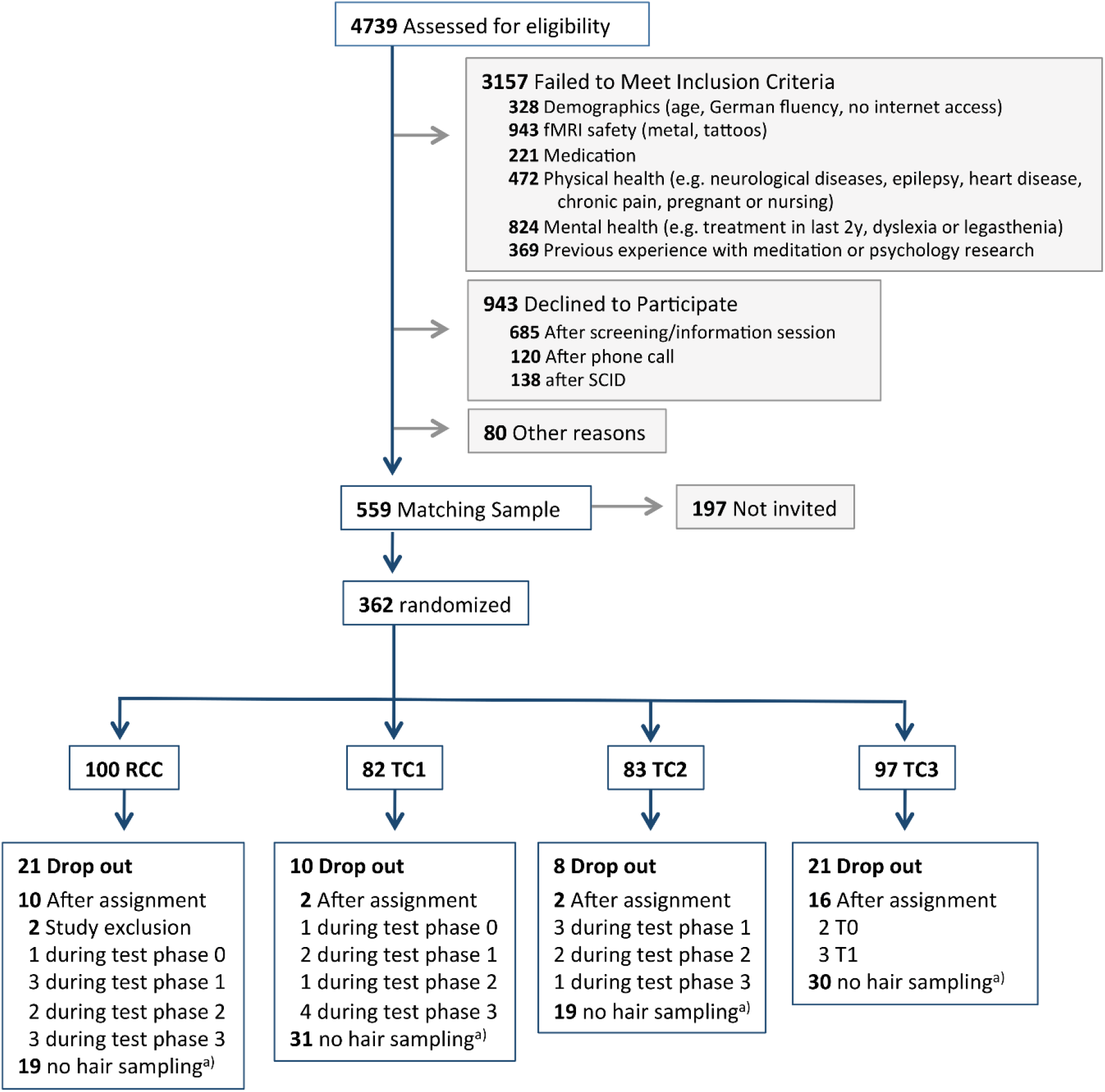
Participant flow chart. This figure combines numbers from two recruitment periods in 2012/2013 and 2013/2014. fMRI denotes functional magnetic resonance imaging; SCID, Structural Clinical Interview for DSM-IV Disorders (Axis I and Axis II); RCC, retest control cohort; and TC, training cohort. Adapted from Singer et al. (2016), chapter 7. ^a)^ Reasons for no hair sampling throughout were boldness or opting out.

### 2.2 Training program

The ReSource Project examined the specific effects of three commonly practiced types of mental training, namely attention-, socio-emotional or socio-cognitive based practices. For this purpose, the training program was parceled into three separate modules (Presence, Affect, and Perspective), each of which cultivated distinct contemplative capacities over three months (Figure 1A; Singer et al., 2016). Every module began with a 3-day retreat during which professional teachers introduced participants to the conceptual core and the relevant practices of a given module. Afterwards, participants attended weekly 2-hour group sessions, and were asked to exercise the respective module’s two core practices for 30 minutes daily on five days per week using a tailor-made app and online platform.

The psychological processes targeted in the Presence module are attention and interoceptive awareness. Its core practices are Breathing Meditation and Body Scan, both of which are classical mindfulness-based exercises also implemented in the MBSR program. The Affect module targets social emotions such as compassion, loving kindness and gratitude. It also aims to enhance prosocial motivation and dealing with difficult emotions. These skills are targeted through the core practices Loving-kindness Meditation and Affect Dyad. In the Perspective module, participants train metacognition and perspective-taking on self and others through the core practices Observing-thoughts Meditation and Perspective Dyad.

The two contemplative dyads are partner exercises that were developed for the ReSource training (Kok and Singer, 2017). They address different skills such as perspective taking on self and others (Perspective Dyad) or gratitude, acceptance of difficult emotions and empathic listening (Affect Dyad) but are similar in structure (for details see also Singer et al., 2016). In each 10-min dyadic practice, two randomly paired participants share their experiences with alternating roles of speaker and listener. The dyadic format is designed to foster interconnectedness by providing opportunities for self-disclosure and non-judgmental listening (Kok & Singer, 2017; Singer et al., 2016).

The distinction between the Affect and Perspective modules reflects research identifying distinct neural routes to social understanding: One socio-affective route including emotions such as empathy and compassion, and one socio-cognitive route including the capacity to mentalize and take perspective on self and others (for details on the scientific backbone of this division see Singer, 2006, 2012).

### 2.3 Study Design

Participants were assigned either to one of two 9-month training cohorts completing all three training modules in different orders (TC1, initial n=80, n for present study=48; and TC2, initial n=81, present n=62), a 3-month Affect only training cohort (TC3, initial n=81, present n=49) or a retest control cohort (RCC, initial n=90, present n=68) (Figure 1B; Singer et al., 2016, chapter 7). Cohort assignment was completed using bootstrapping without replacement to ensure the formation of demographically homogeneous groups. TC1 and TC2 began their training with the attention-based Presence module. Subsequently, they underwent Affect and Perspective training in different orders, thus controlling for sequence effects. TC3 was conducted to isolate the specific effects of the Presence module from the Affect module. The study followed a mixed design, in which most but not all participants received all types of training. Training and data collection took place between April 2013 and February 2016.

### 2.4 Hair cortisol (HC) and hair cortisone (HE) concentrations

HC and HE concentrations are indicative of systemic cortisol exposure and markers of chronic stress (Stalder et al., 2017). Levels of the inactive cortisol metabolite and precursor molecule cortisone have been suggested to yield a complementary, potentially more stable glucocorticoid signal alongside cortisol itself (Stalder et al., 2013; supplement). While the precise mechanism behind HC/HE accumulation is incompletely understood, it is assumed that during hair growth, free cortisol and cortisone molecules are continuously incorporated into follicles, proportional to their overall concentration in the physiological system. HC and HE concentrations in a 1 cm hair segment are thus assumed to indicate the cumulative systemic cortisol/cortisone exposure over an approximately 1 -month period (Stalder, et al., 2017).

For their assessment, hair strands were taken as close as possible to the scalp from a posterior vertex position at T0 and after each training module (at T1, T2 and T3). Hair samples were wrapped in aluminum foil and stored in the dark at room temperature until assay at the Department of Psychology, TU Dresden, Germany. Based on the assumption of an average hair growth rate of 1 cm/month (Wennig, 2000), we analyzed the proximal 3 cm segment of hair to assess cortisol/cortisone accumulation over each 3-month period. Hormone concentrations were measured using liquid chromatography-tandem mass spectrometry (LC-MS/MS), the current gold-standard approach for hair steroid analysis (Gao, Kirschbaum, Grass, & Stalder, 2016), following our previously published protocol with a limit of quantification for cortisol and cortisone below 0.09 pg/mg and intra- and inter-assay CVs between 3.7 and 8.8% (Gao et al., 2013). A first assay of samples collected at baseline was conducted in 2015 to allow researchers to address cross-sectional research questions (Engert et al., 2018) before termination of the longitudinal data collection. Thirty-eight samples were used up in this analysis. For the current longitudinal research aim, the remaining baseline samples were re-assayed jointly with all additional samples (assessed at T1, T2 and T3) to avoid potential systematic effects of storage time and minimize reagent batch effects. Specifically, all samples of one participant were always run with the same reagent batch to avoid intra-individual variance due to batch effects. For the sake of completeness, we also report data on dehydroepiandrosterone (DHEA) to cortisol ratios, which were assessed in the same hair samples, in Supplementary Results B.

### 2.5 Subjective stress measures

Self-reported chronic stress was measured on the basis of the summary score of the Perceived Stress Scale (PSS; Cohen, Kamarck, & Mermelstein, 1983) as well as the global stress score of the Trier Inventory for Chronic Stress (TICS; Schulz & Schlotz, 1999). The 10-item PSS is the most widely used psychological instrument for measuring the perception of stress. It focuses on the degree to which situations in the past month are appraised as unpredictable, uncontrollable and overloaded, and produces one summary stress score. The 39-item TICS captures a time span of 1-3 months and measures six aspects of chronic stress (work overload, worries, social stress, lack of social recognition, work discontent and intrusive memories), and one global stress score. Both questionnaires have satisfactory reliability and validity (Cohen et al., 1983; Schulz & Schlotz, 1999).

### 2.6 Measures of training engagement

To examine causes of individual variability in training effects we assessed two measures of training engagement: practice frequency, objectively traced via our online training platform, and selfreported liking of the different training modules. Details on the measurement and analysis of both metrics are provided in the Supplementary Methods. Practice frequency is a particularly interesting metric as it provides insights to the impact of training dosage.

### 2.7 Statistical analysis

#### Data processing

Raw HC and HE data were each treated with a natural log transformation to remedy skewed distributions. Across the full sample of each dependent measure, any values diverging more than 3 SD from the mean were labeled outliers and winsorized to the respective upper or lower 3 SD boundary. In previous ReSource publications, data has been analyzed as change scores (e.g. Valk et al., 2017). However, change scores can only be computed if a set of consecutive measures is available. Because for HC and HE we had larger than usual dropout rates (Table 1), we chose to analyze the data as simple scores to be able to use all available samples.

**Table 1.**
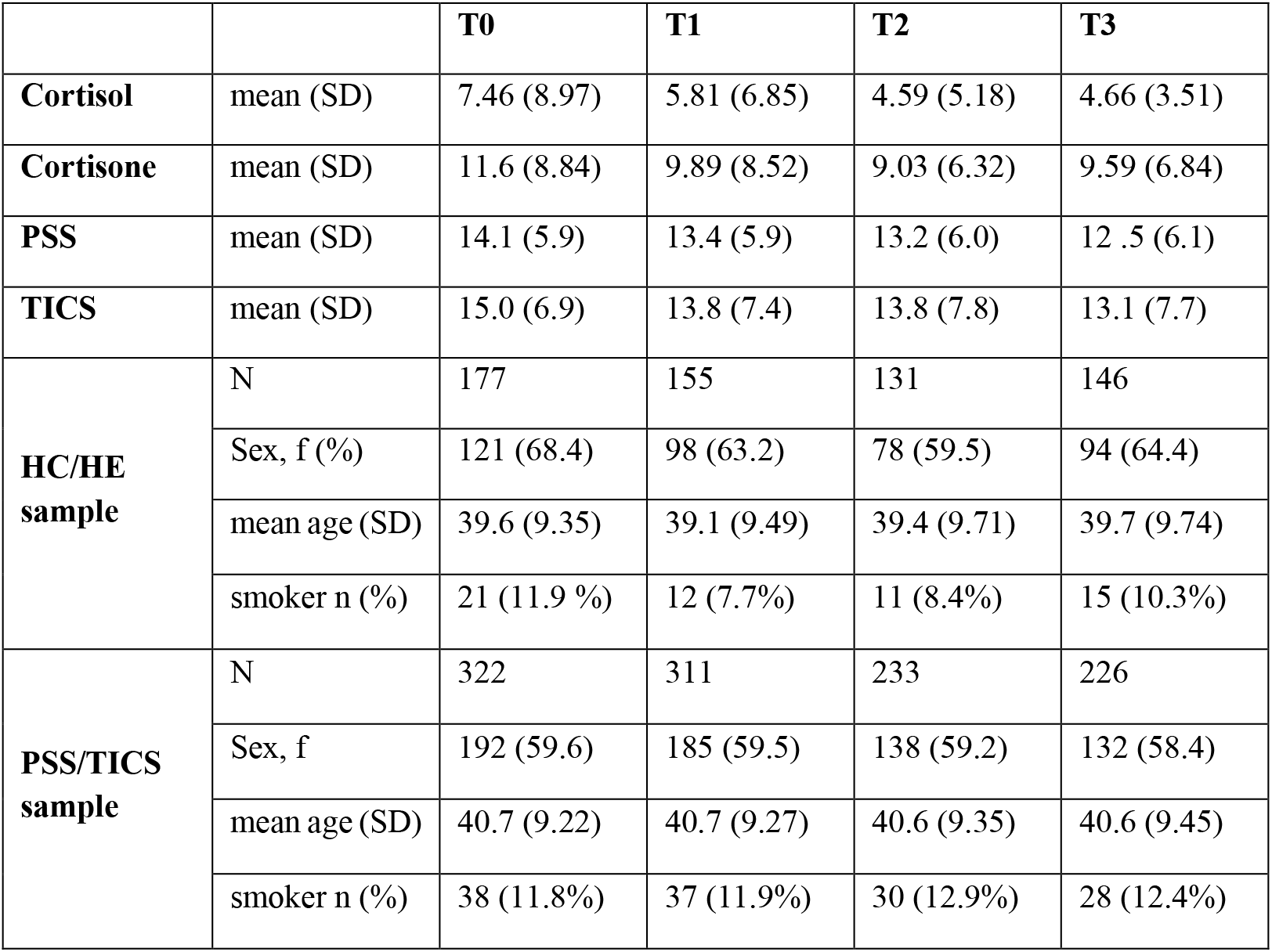
Raw data and demographic characteristics of samples. “HC/HE sample” refers to participants with at least one usable sample of either HC or HE at the given timepoint; “TICS/PSS sample” to participants with one or more self-report rating. More men than women had short hair or were bold, leading to a higher % of women in the HC/HE sample. HC denotes hair cortisol, HE, hair cortisone; PSS, Perceived Stress Scale; TICS, Trier Inventory for Chronic Stress, f, female; m, male; SD, standard deviation. For extensive detail on the demographic characteristics of the sample see Singer et al., 2016. Baseline associations are described in supplementary results A.

|  |  | **T0** | **T1** | **T2** | **T3** |
| --- | --- | --- | --- | --- | --- |
| **Cortisol** | mean (SD) | 7.46 (8.97) | 5.81 (6.85) | 4.59 (5.18) | 4.66 (3.51) |
| **Cortisone** | mean (SD) | 11.6 (8.84) | 9.89 (8.52) | 9.03 (6.32) | 9.59 (6.84) |
| **PSS** | mean (SD) | 14.1 (5.9) | 13.4 (5.9) | 13.2 (6.0) | 12.5 (6.1) |
| **TICS** | mean (SD) | 15.0 (6.9) | 13.8 (7.4) | 13.8 (7.8) | 13.1 (7.7) |
| **HC/HE sample** | N | 177 | 155 | 131 | 146 |
|  | Sex, f (%) | 121 (68.4) | 98 (63.2) | 78 (59.5) | 94 (64.4) |
|  | mean age (SD) | 39.6 (9.35) | 39.1 (9.49) | 39.4 (9.71) | 39.7 (9.74) |
|  | smoker n (%) | 21 (11.9 %) | 12 (7.7%) | 11 (8.4%) | 15 (10.3%) |
| **PSS/TICS sample** | N | 322 | 311 | 233 | 226 |
|  | Sex, f | 192 (59.6) | 185 (59.5) | 138 (59.2) | 132 (58.4) |
|  | mean age (SD) | 40.7 (9.22) | 40.7 (9.27) | 40.6 (9.35) | 40.6 (9.45) |
|  | smoker n (%) | 38 (11.8%) | 37 (11.9%) | 30 (12.9%) | 28 (12.4%) |

#### Significance testing

All statistical analyses were conducted in the statistical software R (version 3.5.1, R Core Team 2018) and with an α-threshold of ≤ 0.05. For completeness, significant and trend-level results (defined as .05 < p <. 10) are reported. Hypotheses were tested by means of multivariate linear mixed models (LMMs), which are robust to unbalanced and incomplete data in longitudinal designs. Models were fit using the function ‘lmer’ of the r package “lme4” (Bates et al., 2015). In models predicting HC or HE, age and sex were included as covariates to account for their potential influence on hormone concentrations (Stalder et al., 2017). The full model included the following terms:

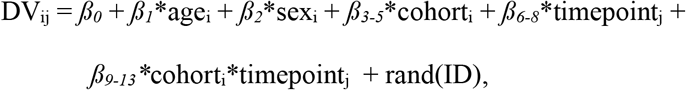

where DV = dependent variable (cortisol, cortisone or subjective stress scores assessed via PSS and TICS), *β0* = intercept, i = subject ID, j = measurement timepoint (T0, T1, T2, T3), rand(ID) = random intercept per subject.

In an omnibus test, we first evaluated whether the respective dependent variable differed as a function of training routine or of time, by testing for an interaction of training by time. Full models with the above outlined terms were compared with reduced models lacking the interaction term via likelihood ratio tests (Dobson, 2002). If TCs differed from the RCC over time, the interaction model will provide a significantly better fit. To ensure accurate model comparisons, models were fitted with the maximum likelihood (ML) method. Effect sizes of significant interactions were calculated as omega squared (ω^2^) by dividing the variance of the residuals of the full model by the variance of residuals of the reduced model and subtracting the outcome from 1 (Xu, 2003). The resulting effect sizes were classified as small (ω^2^≥.010), medium (ω^2^ ≥.059) or large (ω^2^ ≥.138) (Kirk, 1996). Given a significantly better fit of an interaction model, potential differences between training modules and individual measurement timepoints were evaluated in detail by contrasting model estimates through follow-up t-tests, computed through the function ‘lsmeans’ of the package ‘lsmeans’. To this end, models were re-fitted with the restricted maximum likelihood (REML) method to obtain unbiased model estimates. Follow-up contrasts were thus conducted within the LMM framework not corrected for multiple comparisons. The results of model residual checks are reported in the supplementary materials (supplementary results C).

#### Power analysis

Since the present study is part of a large-scale investigation (the ReSource Project) with numerous sub-projects, the sample sizes of the cohorts could not be tailored to this study. To determine whether the analyses planned here were sufficiently powered to be meaningful, we used the function ‘powerSim’ from the package simr to simulate what effect sizes they were sensitive to, given our sample size. Power analyses were based on 1000 runs and conducted in light of our hypotheses, meaning effects were simulated after Presence, Affect, and/or Perspective modules. Depending on the exact pattern of effects, sufficient power was given to detect a minimum of 19-34% change in HC, 11-22% in HE, 11-21% in PSS and 13-21% in TICS as a function of training (supplementary materials, Table S3). While there are no previous studies that may serve as guidelines for reasonable effect sizes regarding HE or HC reduction after mental training, we had previously detected a large relative decreases in acute cortisol reactivity of 32-59% following the same training as employed here (Engert et al., 2017). Our analyses were adequately powered to detect effects even at the lower end of that spectrum.

#### Baseline-matched analysis

In randomized clinical trials, baseline differences are by definition the product of chance rather than representing a latent confound (de Boer et al., 2015). It is, however, possible that participants with higher baseline values are disproportionally assigned to the training cohorts by chance, leading to an overestimation of training effects through conflation with regression to the mean. Following up on our planned analyses, we examined to what extent such a pattern may have influenced the study outcomes. To this end, we selected a subsample of participants with matched baseline characteristics and tested whether our results hold in these data. Similar to clinical studies in which patients are matched to control participants based on their baseline characteristics, we here matched TC participants to RCC participants with respect to their baseline glucocorticoid levels and sex, using the function ‘matchit’ of the R package ‘Match.It’ with replacement (Ho et al., 2011). Each TC was matched separately, with the respective cohort serving as the subject pool from which participants could be drawn multiple times. Participant samples were not artificially duplicated in this process, but instead, the relative matching frequency of each participant was recorded as a weight (higher weights representing multiple matching). Weights for RCC participants were set to 1. Unmatched participants were excluded from the analysis; participants who had missing samples at baseline but provided data at a later timepoint were included. Analysis of HC and HE in the thus generated sample was repeated as for the main analysis, with the addition of a weighting parameter based on the frequency of matching.

## 3. Results

Of 332 participants recruited for the ReSource project, 227 provided samples of hair cortisol or cortisone and 326 provided subjective stress ratings at one or more of the four timepoints of measurement (see Table 1 for samples and demographic characteristics; Figure 2 and Table S1 for sample size and reasons for missingness). Over the nine months of training, HC and HE levels showed high consistency in their pattern of change (Figure 3). A significant cohort by time interaction was detected for HC (χ^2^=30.87, df=7, *p*<.001, ω^2^=0.104) as well as HE (χ^2^=19.14, df=7, *p*=.008, ω^2^= 0.036). Follow-up contrasts (Tables S4 & S5) showed that HC and HE levels remained stable in the no-training RCC. With mental training HC and HE levels decreased steadily until six months into the training regimen, regardless of practice content (Figure 3). At six months, hair glucocorticoid levels in all TCs were significantly reduced compared to the respective pre-training baseline. At the final nine months measurement, HC and HE levels remained stable at this lowered level or regressed slightly towards baseline, but always remained significantly below baseline. Change in HC but not HE concentration was significantly and negatively associated with practice compliance (χ^2^=4.46, *p*=.035, est.: −0.140+/0.066, ω^2^=0.025), suggesting that greater training dosage lead to stronger HC reduction. Neither HC nor HE change was associated with self-reported liking of the modules.

**Figure 3.**
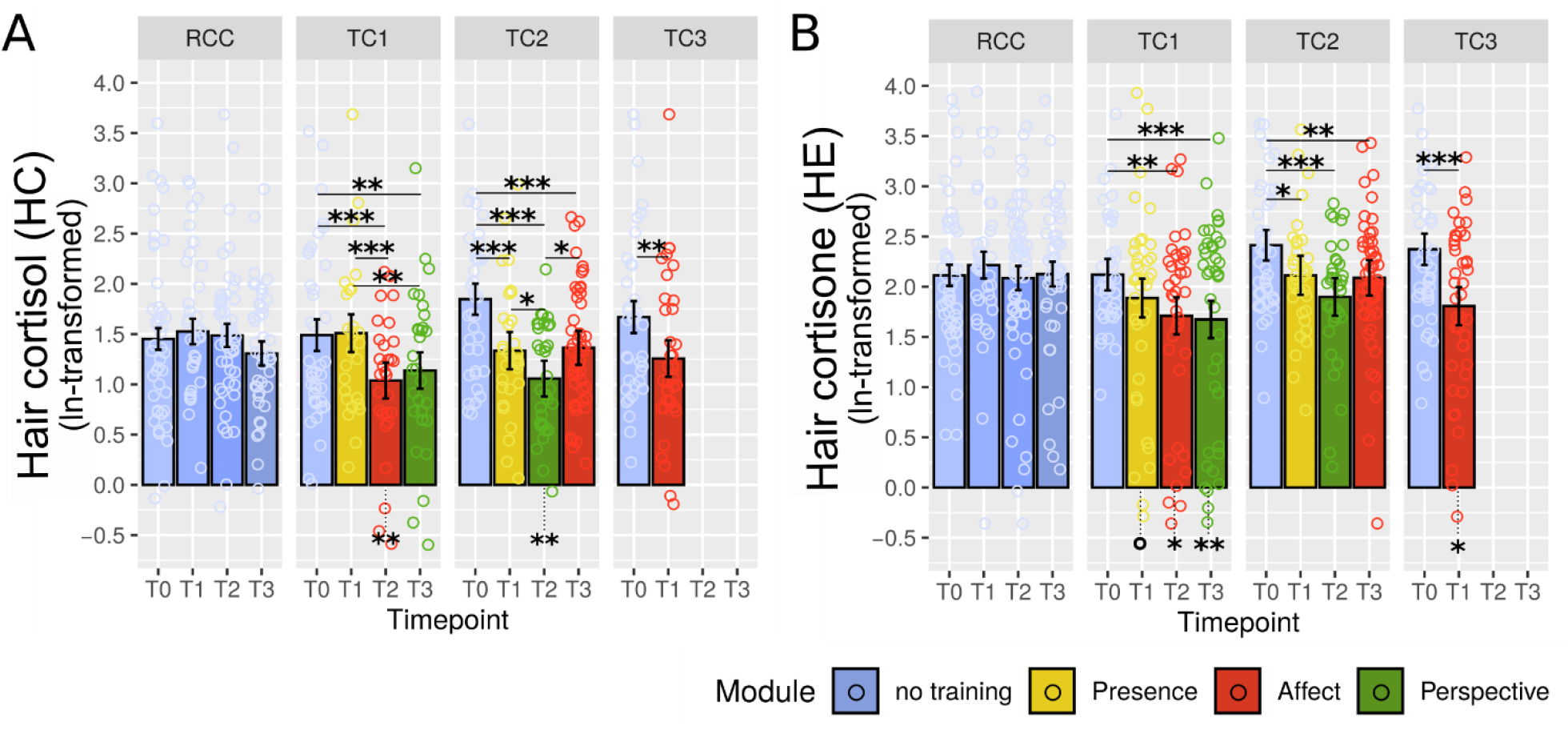
Training effects on cortisol and cortisone levels in hair. Estimated A) hair cortisol and B) hair cortisone levels were derived from the linear mixed model analysis as a function of training cohort and timepoint. Error bars represent +/− 1 SE, each circle represents one data point. Asterisks below bars indicate comparison to RCC. °: trend at 0.05 <*p* <=.1; *: significant at *p* <=.05; **: significant at *p* <=.01; ***: significant at *p* < =.001. See Tables S4 and S5 for a full list of contrast outcomes.

Visualization in Figure 3 suggests that mean HC and HE baseline (T0) values differ somewhat across cohorts, with TC2 and TC3 displaying numerically higher values than TC1 and RCC. In randomized controlled trials, testing for significance of baseline differences is redundant because random subject assignment ensures that any observed baseline differences must arise by chance (de Boer et al., 2015). Nonetheless, an illustrative baseline-matched weighted LMM analysis suggested that results would be comparable in a sample with matched baseline levels (Figure 4). Post-hoc contrasts revealed a similar pattern as in the main analysis. Reduced significance overall indicates a potential overestimation of training effects due to skewed baselines but may also be attributable to the reduced sample size of this additional control analysis, in which several TC participants with relatively higher baselines were excluded.

**Figure 4.**
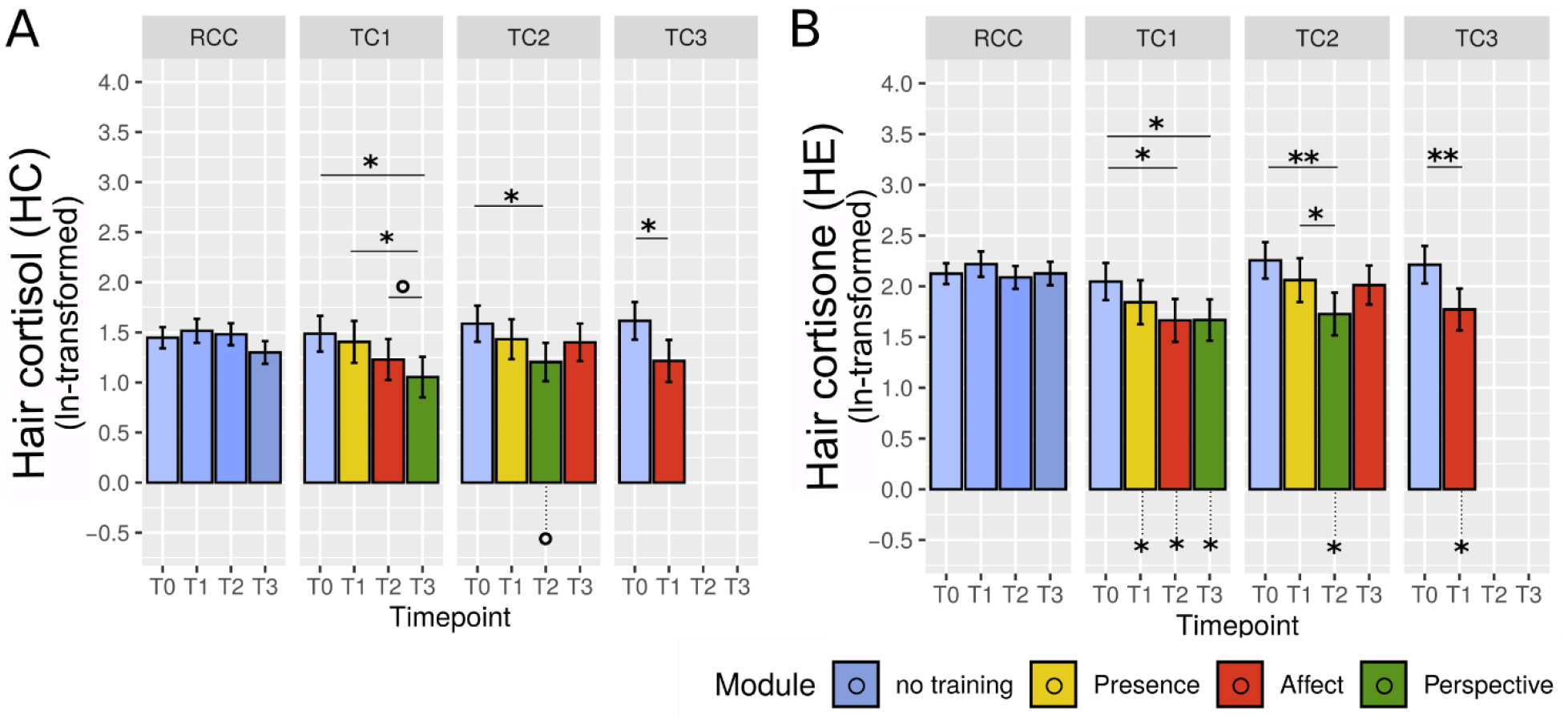
Training effects in baseline-matched analysis. Estimated A) hair cortisol and B) hair cortisone levels were derived from LMM analysis in a sample of participants with matched baseline HE and HE levels across cohorts, generated on the basis of the study participant pool. Participants from each TC were matched to RCC participants with replacement depending on their baseline glucocorticoid levels and gender. Error bars represent +/− 1 SE, each circle represents one data point. Asterisks below bars indicate comparison to RCC. °: trend at 0.05 <*p* <=.1; *: significant at *p* <=.05; **: significant at *p* <=.01; ***: significant at *p* <=.001.

In another analysis of potential bias, baseline HC and HE levels did not differ between TC participants who dropped out from hair sampling during the study and those who did not (HC: *t*(112)=0.5, *p*=.62; HE: *t*(125)=-0.7, *p*=.49), demonstrating that there was no selective drop-out.

In the analysis of subjective-psychological stress reduction, the cohort by time interaction was significant for PSS (χ^2^=22.20, df=7, *p*=.002, ω^2^=0.030) but only marginal for TICS values (χ^2^=13.66, df=7, *p*=.058, ω^2^=0.018) (Figure 5). Follow-up contrasts of PSS scores suggested that participants reported lowest subjective stress experience following the Perspective module, but only in TC1. Exploratory LMM analyses of all samples showed no significant association between PSS or TICS scores with HC or HE concentrations throughout the study.

**Figure 5.**
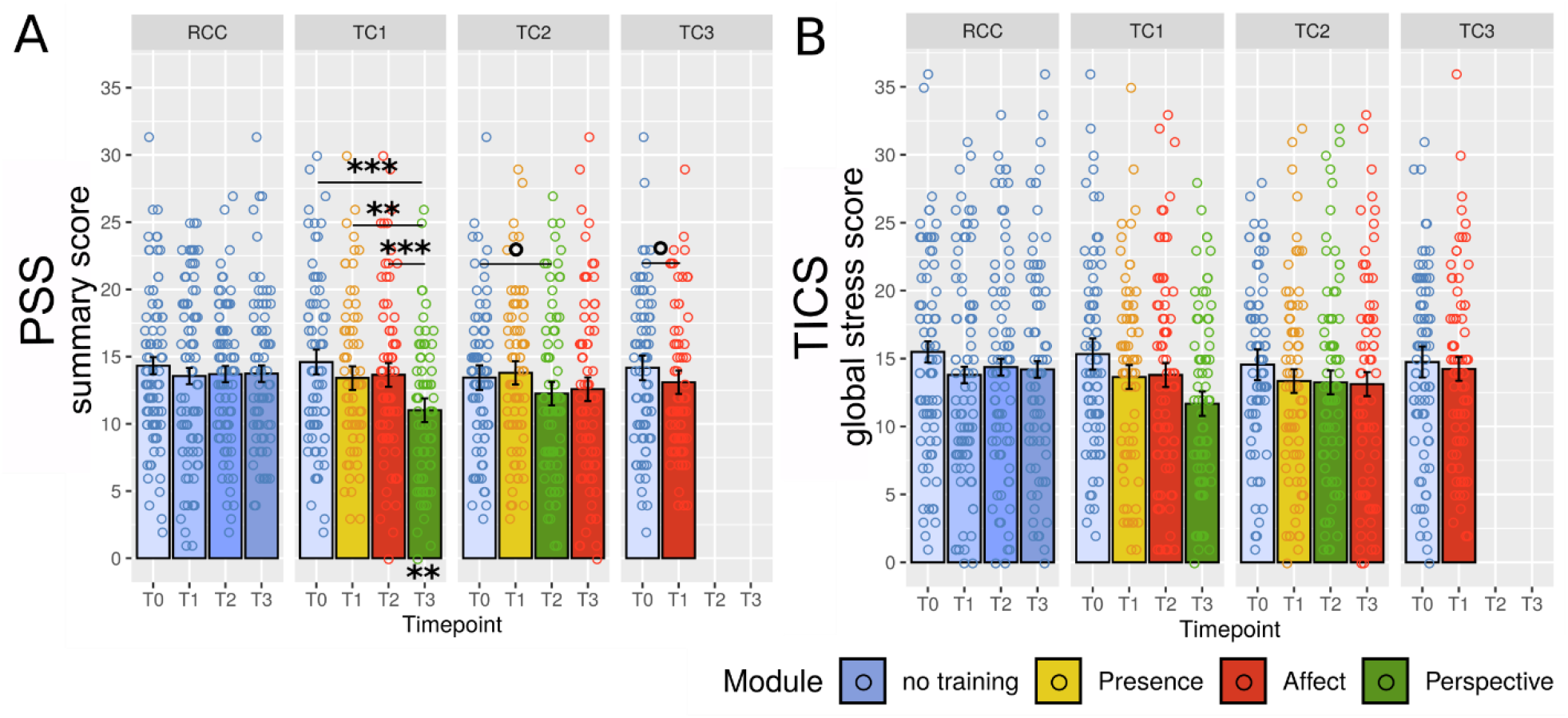
Training effects on self-reported long-term stress. Estimated scores of A) Perceived Stress Scale (PSS; Cohen et al., 1983) and B) Trier Inventory for Chronic Stress (TICS; Schulz & Schlotz, 1999) derived from the linear mixed model analysis as a function of training cohort and timepoint. Error bars represent +/− 1 SE, each circle represents one data point. °: trend at 0.05 < p <=.1; *: significant at p <=.05; **: significant at p <=.01; ***: significant at p <=.001.

Similar to HC, PSS change was negatively associated with practice frequency (χ^2^ =4.99, *p*=.025, est.:-0.591+/-0.264, ω^2^=0.010), and additionally with liking of the modules (χ^2^ =9.34, *p*=.002, est.: −0.975+/-0.318, ω^2^=0.019; see also supplementary methods). However, the effect of practice duration disappeared when controlling for participants’ ratings of how much they liked the respective module, suggesting that module enjoyment was the latent driver of the practice association. The effect of liking contrarily persisted even when controlling for practice frequency (χ =6.07, *p*=.014, est.:-0.815+/-0.330, ω^2^=0.012). Considering that change in HC was not associated with self-reported liking, measures of stress and training engagement appear to cluster in subjective self-report measures, perhaps reflecting the lack of psychoendocrine covariance that is commonly reported in the stress literature (Campbell & Ehlert, 2012; Engert et al., 2018)

## 4. Discussion

The present investigation examined whether the 9-month-long training of different types of contemplative mental practice affects physiological indices of chronic stress. Our results show that daily mental training over 3-6 months can buffer the long-term systemic stress load of healthy adults, reflected in a reduction of cortisol and cortisone accumulation in hair. This effect was independent of specific training content, positively associated with practice frequency for HC, and reached a ceiling after six months of training. At the same time, consistent significant difference to baseline was only achieved after six months of training, suggesting that reliable long-term benefits in HPA axis activity emerge only after a relatively long period of intense training. This may explain why previous studies found no HC reductions after the typical 8-12 weeks of mindfulness-based training (e.g. Gotink et al., 2017; Nery et al., 2019), with the exception of one pilot study with 18 smokers (Goldberg et al., 2014).

In an earlier ReSource Project publication with the same participant sample (Engert et al., 2017), we found that Affect and Perspective training selectively reduced acute salivary cortisol release in response to a stressful psychosocial laboratory challenge, the Trier Social Tress Test (TSST; Kirschbaum et al., 1993). This differentiated pattern of results between indices of acute compared to chronic HPA axis activity suggests that distinct processes may underlie change in either type of activity. It is conceivable that stress “immunization” to a psychosocial challenge is best achieved with a training that targets social processes, such as the dyadic partner exercises implemented in the Affect and Perspective modules. In contrast, the cumulative HPA axis activity as monitored in hair may reflect the more low-grade and continuous strain inherent to various daily hassles (Almeida, 2005; DeLongis et al., 1982; Lazarus & Folkman, 1984), which appears to be equally buffered by all three mental training techniques. While in the ReSource Project, we find differential training effects of the three realized practice types on many levels of observation (Singer & Engert, 2019), some changes appear to need time to develop, irrespective of practice type (see also Bornemann & Singer, 2017).

Changes in self-reported measures of chronic stress were unrelated to changes in HC and HE. This lack of psychoendocrine covariance is a recurring phenomenon in stress research (e.g. Campbell & Ehlert, 2012; Engert et al., 2018) and may be particularly pronounced in retrospective assessments, where the specific dynamics of different stress systems become washed out (Schlotz et al., 2008). While we expected a pronounced reduction in subjective stress, perhaps exaggerated through biases, change in self-report measures was inconsistent and did not show the robust reductions reported in previous studies (e.g. Khoury et al., 2015). It is possible that participants experienced the uniquely large-scale testing of the ReSource Project as straining, leading to this discrepancy. To this effect, we previously found that the realized training practices can also be experienced as effortful (Lumma et al., 2015).

Despite our large number of participants, the number of dropouts from the hair glucocorticoid assessment – partly attributable to the optional nature of this assessment – is a limitation of the current work. Importantly, however, most participants dropped out of this assessment already at study baseline and quality control analyses revealed no evidence for systematic dropouts. For the overall interpretation of this work it should be noted that cumulative indices of HPA-axis regulation like HC and HE do not allow specific conclusions about the physiological mechanisms leading to cortisol or cortisone levels in hair. Changes in diurnal cortisol dynamics, cortisol release under acute stress or under more low-level strains may all contribute to lower HC/HE levels.

In sum, the present investigation provides evidence that mental training has a beneficial effect on individuals’ long-term physiological stress load, irrespective of specific practice type. With HC and HE, we targeted the cumulative burden of frequent HPA axis activation, which is particularly maladaptive and related to ill-health. Our results thus point to one mechanism via which mental training can exert positive effects on practitioners’ health status in general: By lowering systemic cortisol exposure, regular practice of about 30 minutes daily for six months may reduce vulnerability for stress-associated disease. We conclude that to achieve chronic stress reduction at the level of HPA axis activation, it is worth to practice more, and to carry on mental practice beyond the typical eight-week training period of mindfulness-based stress reduction programs currently offered in Western societies.

## Data availability

In line with the GDPR, our data cannot be shared publicly because we did not obtain explicit participant agreement for data-sharing with parties outside the MPI CBS. Original data are available upon request (contact via).

## Author contributions

T. Singer. initiated and developed the ReSource Project and secured all funding except for the hair glucocorticoid analysis. T. Singer and V.E. designed the experiment. V.E., L.P., P.V. and R.L. were involved in data curation; L.P. and P.V. analyzed the data. C.K. funded the hair analyses. C.K. and T. Stalder were responsible for planning, performing and interpreting the hair glucocorticoid analysis. V.E. and L.P. drafted, and all authors critically revised the manuscript.

