## Supplementary materials for "Contemplative mental training reduces hair glucocorticoid levels in a randomized clinical trial"

### *Supplementary Methods*: Practice frequency and liking of training

*Data collection and processing.* After completing each module, participants rated how much they liked practicing the two core exercises of the respective module on a 5-point Likert scale from 1 (“not at all”) to 5 (“very much”) (see also Singer et al. 2016; Appendix H). Practice frequency was tracked online through a custom-made ReSource training platform and is here reported as the average number of times participants practiced each exercise per week. To limit the number of exploratory analyses and to improve variable stability, we averaged liking and practice scores across the two core exercises of each module.

By nature, liking and practice scores were only available from training cohort (TC) participants, and not collected at baseline (T0). To be able to model change in our dependent variables directly as a function of liking and practice scores, we thus had to generate change scores. These were calculated by taking the difference between raw measurements from each set of consecutive timepoints (T1-T0, T2-T1 and T3-T2). HC and HE change scores were calculated from the ln-transformed data; PSS and TICS change scores were calculated from the raw summary scores (for available samples and raw data used in the exploratory analyses see supplements, Table S2). Outliers in the data were identified and winsorized following the same procedure as described in the main methods section.

*Significance testing.* In line with our main analyses, we used linear mixed models (LMMs) and full-to-reduced model comparisons to assess whether the addition of liking or practice scores to the model significantly improved explained variance in HC, HE, PSS, or TICS change. We additionally assessed the influence of an interaction term, liking x module, to examine whether liking of the Perspective module in particular was implicated in the reduction of subjective stress. In the latter analysis, we combined data from the different training cohorts and grouped them by training module.

***Supplementary Results A*: Baseline analyses.**

At baseline (T0), age significantly positively correlated with HC (Pearson $r$ = 0.165, $p$ = .039), and positively but non-significantly with HE levels (Pearson $r$ = 0.123, $p$ = 0.102). T-tests revealed no significant difference between males and females in HC or HE levels. Because the literature suggests an influence of sex on HC (Stalder et al., 2017), we nonetheless controlled for both sex and age in our models. Baseline exploratory analyses showed that concentrations of HC and HE were highly positively correlated ($r$=0.608, $p$<.001). The same was true for T0 questionnaire scores of PSS and TICS ($r$=0.692, $p$<.001). HC and HE were not related to PSS scores at baseline, but, surprisingly, significantly negatively associated with baseline TICS scores (HC: $r$=-0.204, $p$=.011; HE: $r$=-0.151, $p$=.046).

***Supplementary Results B*: Dehydroepiandrosterone to cortisol ratios**

In addition to hair cortisol and cortisone, the ratio of hair cortisol (HC) to dehydroepiandrosterone (DHEA) is understood to be indicative of hypothalamic-pituitary-adrenal functioning, and is suggested to account for the neurotrophic effects of each hormone (Maninger et al., 2009). Since there is little published data on DHEA to cortisol ratios measured in hair (DHEA/HC), and for the sake of completeness, we also report results relating to the latter.

DHEA was assayed in the hair using the same methods as for HC and HE and reported in pg/mg. DHEA/HC ratios were calculated by dividing raw HC measures by raw DHEA measures. The resulting values were ln-transformed and treated for outliers as described in the main methods section to achieve a normal distribution. At baseline, DHEA/HC ratios correlated positively with age ($r$=0.271, $p$=.001), HE ($r$=0.360, $p$ < .001) and HC ($r$=0.768, $p$<.001), and negatively with DHEA ($r$=-0.658, $p$<.001). T-test showed that women had significantly higher DHEA/HC ratios (mean [SD] = 1.58 [1.13]) than men (mean [SD] = 0.98 [0.98]) ($t$(141)=3.13, $p$=.002).

Subsequently, potential effects of training were evaluated using the same statistical approach as for the main analyses in the paper. Full to reduced model comparison showed a significant effect of the cohort x time interaction term on DHEA/HC ratios ([illegible]=23.17, df=7, $p$=.002, $\omega^2$=0.080). Like the pattern observed in HC and HE, DHEA/HC ratios appeared stable in the RCC and showed decreases in TC2 and TC3. DHEA/HC ratios of the TC1, however, did not decrease. Post-hoc test showed that participants of the TC3 had significantly lower DHEA/HC ratios at T1 following the Affect training than at T0 ($t$(334)=3.14, $p$=0.010, est.=-0.474+/-0.15). TC2 showed reductions in DHEA/HC ratios at T1 following Presence training and further at T2 following Perspective training, at which point values became significantly lower than at baseline ($t$(317)=3.85, $p$<.001, est.=-0.591+/-0.15). Subsequently, DHEA/HC values increased again towards baseline, such that the difference between T2 and T3 scores trended towards significance ($t$(300)=2.58, $p$=.051, est.=0.355+/-0.14). The results are shown in the Supplementary Figure S2.

***Figure S1.*** **DHEA to hair cortisol (HC) ratios by training cohort and timepoint.**

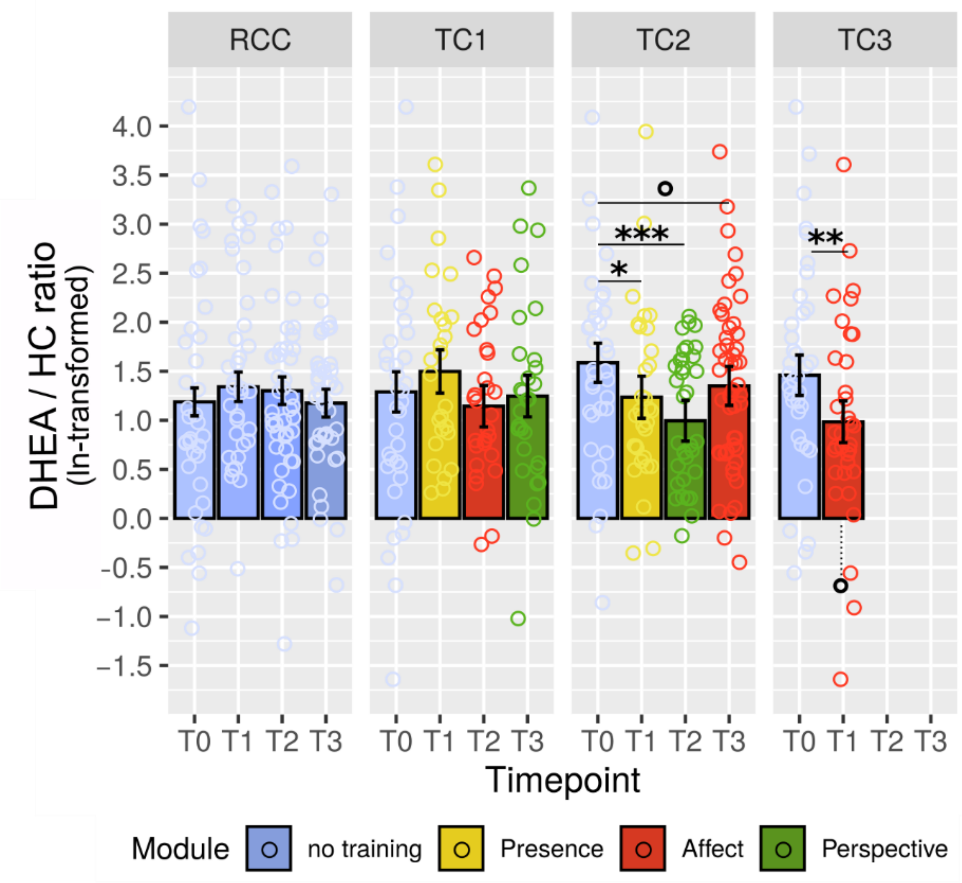

*Note:* Estimated DHEA to hair cortisol (DHEA/HC) ratios were derived from the linear mixed model analysis outlined in supplementary analysis B. Error bars represent 95% confidence intervals, each circle represents one data point. DHEA, dehydroepiandrosterone. °: trend at 0.05 < p <=.1; *: sign. at p <=.05; **: sign. at p <=.01; ***: sign. at p <=.001

***Supplementary Results C*: Model residual checks.**

All models' residuals displayed satisfactory approximation to normal distribution, with the exception of the analysis of HE change data in relation to practice frequency, in which residual distribution had unusually light tails owing to the high kurtosis in HC change. Variance inflation factors of all models' main effects indicated uncritical levels of multicollinearity (Hair Jr., Black, Babin, & Anderson, 1998). Estimates of cook's distances showed no evidence of highly influential observations in HC or HE models, but 1 and 3 influential case(s) in models of PSS and TICS, respectively (indicated through cook's $d > 1$ and/or visual inspection; Fox, 1991). Re-calculating models and contrast estimates after removal of these cases did not alter any of the previous results.

***Table S1*. Availability of raw data and reasons for missing cases**

| Variable | N | Reasons for missingness |
|---|---|---|
| **Age** | 332 | N/A |
| **Sex** | 332 | N/A |
| **HC** | **T0**<br>*Available:* 179<br>*Usable:* 156 | Study dropout (n = 4)<br>Study exclusion (n = 2)<br>No hair sampling throughout (n = 99)[a]<br>No hair sample (missing) (n = 48)<br>< dl (n = 23) |
| | **T1**<br>*Available:* 157<br>*Usable:* 130 | Study dropout (n = 11)<br>No hair sample (missing) (n = 58)<br>< dl (n = 27) |
| | **T2**<br>*Available:* 136<br>*Usable:* 112 | Study dropout (n = 5)<br>TC3 Study completed (n = 49)[b]<br>No hair sample (missing) (n = 25)<br>< dl (n = 24) |
| | **T3**<br>*Available:* 150<br>*Usable:* 124 | Study dropout (n = 8)<br>No hair sample (missing) (n = 3)<br>< dl (n = 26) |
| **HE** | **T0**<br>*Available:* 179<br>*Usable:* 177 | Study dropout (n = 4)<br>Study exclusion (n = 2)<br>No hair sampling throughout (n = 99)[a]<br>No hair sample (missing) (n = 48)<br>< dl (n = 2) |
| | **T1**<br>*Available:* 157<br>*Usable:* 155 | Study dropout (n = 11)<br>No hair sample (missing) (n = 58)<br>< dl (n = 2) |
| | **T2**<br>*Available:* 136<br>*Usable:* 131 | Study dropout (n = 5)<br>TC3 Study completed (n = 49)[b]<br>No hair sample (missing) (n = 25)<br>< dl (n = 5) |
| | **T3**<br>*Available:* 150<br>*Usable:* 146 | Study dropout (N = 8)<br>No hair sample (missing) (n = 3)<br>< dl (n = 4) |
| **TICS, PSS** | **T0**<br>N = 322 | Study dropout (n = 4)<br>Study exclusion (n = 2)<br>No questionnaire data (missing) (n = 4) |
| | **T1**<br>N = 311 | Study dropout (n = 11)<br>No questionnaire data (missing) (n = 4) |
| | **T2**<br>N = 232 (TICS)<br>N = 233 (PSS) | Study dropout (n = 5)<br>TC3 Study completed (n = 76)[b]<br>No questionnaire data (missing) (TICS: n =2, PSS: n = 1) |
| **Practice frequency**[c] | **T0 to T1**<br>N = 225 | **Only assessed in TCs (N = 242)**<br>TCs Study dropout (n = 11)<br>TCs Study exclusion (n = 0)<br>No practice data (missing) (n = 3) |
| | **T1 to T2**<br>N = 149 | TCs Study dropout (n = 3)<br>TC3 Study completed (n = 76)[b]<br>No practice data (missing) (n = 3) |
| | **T2 to T3** | TCs Study dropout (n = 5) |

| | | |
|---|---|---|
| | N = 144 | No practice data (missing) (n = 3) |
| **Liking**[c] | **T0 to T1**<br>N = 217 | **Only assessed in TCs (N = 242)**<br>TCs Study dropout (n = 11)<br>TCs Study exclusion (n = 0)<br>No liking ratings (missing) (n = 14) |
| | **T1 to T2**<br>N = 144 | TCs Study dropout (N = 3)<br>TC3 Study completed (N = 76)[b]<br>No liking ratings (missing) (N = 8) |
| | **T2 to T3**<br>N = 139 | TCs Study dropout (N = 5)<br>No liking ratings (missing) (N = 8) |

*Notes:* dl: detection limit; HC: hair cortisol; HE: hair cortisone; PSS: Perceived Stress Scale (Cohen, S., Kamarck, T. & Mermelstein, 1983); TICS: Trier Inventory for Chronic Stress (Schulz & Schlotz, 1999).

[a] Reasons for no hair sampling throughout were boldness or opting out

[b] from the total 81 TC3 participants, n = 5 were study dropouts before T2 and an additional n = 27 had no hair sampling throughout

[c] not including data from dropouts of the present study (i.e. participants without HC, HE, TICS and PSS data)

***Table S2.*** **Descriptives of change scores used in exploratory analyses.**

| | | **T0 to T1** | **T1 to T2** | **T2 to T3** |
|---|---|---|---|---|
| **HC** | sample n | 98 | 103 | 92 |
| | mean (SD) | -0.286 (0.83) | -0.196 (0.80) | -0.002 (0.62) |
| **HE** | sample n | 126 | 132 | 118 |
| | mean (SD) | -0.247 (0.90) | -0.138 (0.96) | 0.062 (0.85) |
| **PSS** | sample n | 309 | 271 | 225 |
| | mean (SD) | -0.616 (5.21) | -0.193 (5.33) | -0.71 (5.33) |
| **TICS** | sample n | 309 | 270 | 224 |
| | mean (SD) | -1.29 (5.20) | 0.118 (5.21) | -0.747 (5.05) |
| **practice**[a] | sample n | 225 | 149 | 144 |
| (n/week) | mean (SD) | 4.38 (1.09) | 3.80 (0.77) | 3.51 (0.84) |
| **liking**[a] | sample n | 217 | 144 | 139 |
| (rating) | mean (SD) | 3.99 (0.68) | 3.60 (0.83) | 3.72 (0.83) |

*Notes:* Hair cortisol (HC) and hair cortisone (HE) change scores were calculated from the ln-transformed data. T0 to T1 etc. refer to the change intervals between two timepoints of data sampling (see Figure 1 B, study design). PSS: Perceived Stress Scale (Cohen, Kamarch, & Mermelstein, 1983); TICS: Trier Inventory for Chronic Stress (Schulz & Schlotz, 1999).

[a] not including data from dropouts of the present study (i.e. participants without HC, HE, TICS and PSS data)

***Table S3.* Results of power analysis**

| **Effects simulated** | **HC**[a] (min. *ß*s) | **HE**[a] (min. *ßs)* | **PSS**[b] (min. *ßs)* | **TICS**[b] (min. *ßs)* |
|---|---|---|---|---|
| DV reduced after **Pres.** | -0.45 (TC1&TC2)<br>-0.5 (TC1/TC2) | -0.4 (TC1&TC2)<br>-0.45 (TC1/TC2) | -2.0 (TC1&TC2)<br>-2.4 (TC1/TC2) | -2.1 (TC1&TC2)<br>-2.4 (TC1/TC2) |
| DV reduced after **Affect** | -0.3 (all TCs)<br>-0.35 (TC1&TC2)<br>-0.55 (TC1/TC2)<br>-0.6 (TC3) | -0.25 (all TCs)<br>-0.30 (TC1&TC2)<br>-0.45 (TC1/TC2)<br>-0.50 (TC3) | -1.4 (all TCs)<br>-1.6 (TC1&TC2)<br>-2.4 (TC1/TC2)<br>-2.7 (TC3) | -1.5 (all TCs)<br>-1.7 (TC1&TC2)<br>-2.5 (TC1/TC2)<br>-2.8 (TC3) |
| DV reduced after **Pres. & Affect** | -0.3 (all TCs)<br>-0.3 (TC1&TC2)<br>-0.5 (TC1/TC2) | -0.25 (all TCs)<br>-0.25 (TC1&TC2)<br>-0.45 (TC1/TC2) | -1.4 (all TCs)<br>-1.5 (TC1&TC2)<br>-2.1 (TC1/TC2) | -1.5 (all TCs)<br>-1.4 (TC1&TC2)<br>-2.1 (TC1/TC2) |
| DV reduced after **Pres., Affect & Persp.** | -0.45 (all TCs)<br>-0.45 (TC1&TC2)<br>-0.5 (TC1/TC2) | -0.45 (all TCs)<br>-0.40 (TC1&TC2)<br>-0.45 (TC1/TC2) | -2.3 (all TCs)<br>-2.1 (TC1&TC2)<br>-2.4 (TC1/TC2) | -2.4 (all TCs)<br>-2.2 (TC1&TC2)<br>-2.3 (TC1/TC2) |

[a] approximated in steps of 0.05

[b] approximated in steps of 0.1

*Notes:* For four possible effects of training, we calculated approximate minimum *ß* sizes required to achieve at least 80% power to detect a significant group by time interaction in the planned linear mixed models. Displayed are *ß*s for varying consistencies of the effects across training cohorts. Power was estimated based on simulations run 1000 times, and assuming no group differences at baseline and no effect of time. Lowest and highest *ß* values correspond to the following percentage change (relative to mean baseline values): HC, 0.3: 19%, 0.6: 37%; HE, 0.25: 11%; 0.5: 22%; PSS, 1.4: 10%, 2.7: 19%; TICS, 1.5: 10%, 2.8: 19%. HC denotes hair cortisol; HE, hair cortisone; PSS, perceived stress scale; TICS, Trier inventory for chronic stress; TC, training cohort.

***Table S4*. Follow-up contrasts within LMM of HC levels**

| Contrast | Estimate | SE | df | t-value | p-value |
|---|---|---|---|---|---|
| **Against RCC, T1** | | | | | |
| RCC - TC1 | 0.017 | 0.176 | 472 | 0.097 | 0.923 |
| RCC - TC2 | 0.191 | 0.166 | 481 | 1.149 | 0.251 |
| RCC - TC3 | 0.270 | 0.166 | 445 | 1.621 | 0.106 |
| **Against RCC, T2** | | | | | |
| **RCC - TC1** | **0.449** | **0.163** | **435** | **2.748** | **0.006** |
| **RCC - TC2** | **0.429** | **0.155** | **453** | **2.766** | **0.006** |
| **Against RCC, T3** | | | | | |
| RCC - TC1 | 0.169 | 0.167 | 447 | 1.010 | 0.313 |
| RCC - TC2 | -0.057 | 0.146 | 419 | -0.389 | 0.697 |
| **Within RCC** | | | | | |
| T0 - T1 | -0.074 | 0.127 | 340 | -0.586 | 0.559 |
| T0 - T2 | -0.035 | 0.116 | 351 | -0.299 | 0.765 |
| T0 - T3 | 0.145 | 0.120 | 345 | 1.213 | 0.226 |
| T1 - T2 | 0.040 | 0.122 | 341 | 0.324 | 0.746 |
| T1 - T3 | 0.219 | 0.126 | 340 | 1.734 | 0.084 |
| T2 - T3 | 0.180 | 0.114 | 339 | 1.582 | 0.115 |
| **Within TC1** | | | | | |
| T0 - T1 | -0.019 | 0.139 | 352 | -0.136 | 0.892 |
| **T0 - T2** | **0.453** | **0.135** | **353** | **3.342** | **0.001** |
| **T0 - T3** | **0.352** | **0.136** | **355** | **2.591** | **0.010** |
| **T1 - T2** | **0.472** | **0.142** | **317** | **3.313** | **0.001** |
| **T1 - T3** | **0.371** | **0.144** | **323** | **2.581** | **0.010** |
| T2 - T3 | -0.101 | 0.141 | 327 | -0.711 | 0.478 |
| **Within TC2** | | | | | |
| **T0 - T1** | **0.513** | **0.135** | **360** | **3.803** | **<.001** |
| **T0 - T2** | **0.791** | **0.135** | **361** | **5.863** | **<.001** |
| **T0 - T3** | **0.484** | **0.119** | **363** | **4.060** | **<.001** |
| **T1 - T2** | **0.278** | **0.130** | **313** | **2.131** | **0.034** |
| T1 - T3 | -0.028 | 0.122 | 339 | -0.232 | 0.817 |
| **T2 - T3** | **-0.306** | **0.122** | **336** | **-2.514** | **0.012** |
| **Within TC3** | | | | | |
| **T0 - T1** | **0.413** | **0.130** | **368** | **3.178** | **0.002** |

*Notes*: Significant contrasts are highlighted in bold. Omitted are contrasts at baseline and across TCs, which to not address any of the study hypotheses. LMM denotes linear mixed model; HC, hair cortisol; SE, standard error; df, degrees of freedom; RCC, retest control cohort; TC1-3, training cohort 1-3.

***Table S5.* Follow-up contrasts within LMM of HE levels**

| Contrast | Estimate | SE | df | t-value | p-value |
|---|---|---|---|---|---|
| **Against RCC, T1** | | | | | |
| RCC - TC1 | 0.328 | 0.169 | 551 | 1.943 | 0.053 |
| RCC - TC2 | 0.102 | 0.168 | 572 | 0.608 | 0.544 |
| **RCC - TC3** | **0.408** | **0.165** | **545** | **2.479** | **0.013** |
| **Against RCC, T2** | | | | | |
| **RCC - TC1** | **0.376** | **0.157** | **516** | **2.396** | **0.017** |
| RCC - TC2 | 0.187 | 0.156 | 548 | 1.201 | 0.230 |
| **Against RCC, T3** | | | | | |
| **RCC - TC1** | **0.452** | **0.158** | **521** | **2.853** | **0.005** |
| RCC - TC2 | 0.038 | 0.144 | 514 | 0.260 | 0.795 |
| **Within RCC** | | | | | |
| T0 - T1 | -0.101 | 0.133 | 430 | -0.761 | 0.447 |
| T0 - T2 | 0.028 | 0.12 | 448 | 0.232 | 0.817 |
| T0 - T3 | -0.012 | 0.124 | 438 | -0.099 | 0.921 |
| T1 - T2 | 0.129 | 0.129 | 428 | 1.001 | 0.317 |
| T1 - T3 | 0.089 | 0.132 | 424 | 0.673 | 0.501 |
| T2 - T3 | -0.04 | 0.118 | 422 | -0.341 | 0.731 |
| **Within TC1** | | | | | |
| T0 - T1 | 0.233 | 0.139 | 436 | 1.67 | 0.096 |
| **T0 - T2** | **0.41** | **0.141** | **438** | **2.915** | **0.004** |
| **T0 - T3** | **0.446** | **0.138** | **435** | **3.229** | **0.001** |
| T1 - T2 | 0.177 | 0.139 | 396 | 1.275 | 0.203 |
| T1 - T3 | 0.213 | 0.138 | 406 | 1.544 | 0.123 |
| T2 - T3 | 0.036 | 0.139 | 404 | 0.261 | 0.794 |
| **Within TC2** | | | | | |
| **T0 - T1** | **0.3** | **0.142** | **458** | **2.117** | **0.035** |
| **T0 - T2** | **0.515** | **0.143** | **460** | **3.598** | **<.001** |
| **T0 - T3** | **0.325** | **0.125** | **442** | **2.591** | **0.010** |
| T1 - T2 | 0.215 | 0.141 | 396 | 1.524 | 0.128 |
| T1 - T3 | 0.025 | 0.132 | 438 | 0.187 | 0.852 |
| T2 - T3 | -0.19 | 0.133 | 436 | -1.428 | 0.154 |
| **Within TC3** | | | | | |
| **T0 - T1** | **0.566** | **0.135** | **446** | **4.18** | **<.001** |

*Notes*: Significant contrasts are highlighted in bold. Omitted are contrasts at baseline and across TCs, which to not address any of the study hypotheses. LMM denotes linear mixed model; HC, hair cortisol; SE, standard error; df, degrees of freedom; RCC, retest control cohort; TC1-3, training cohort 1-3.
